# Decoding brain functional connectivity implicated in AD and MCI

**DOI:** 10.1101/697003

**Authors:** Sukrit Gupta, Yi Hao Chan, Jagath C. Rajapakse, the Alzheimers Disease Neuroimaging Initiative

## Abstract

Deep neural networks have been demonstrated to extract high level features from neuroimaging data when classifying brain states. Identifying salient features characterizing brain states further refines the focus of clinicians and allows design of better diagnostic systems. We demonstrate this while performing classification of resting-state functional magnetic resonance imaging (fMRI) scans of patients suffering from Alzheimer’s Disease (AD) and Mild Cognitive Impairment (MCI), and Cognitively Normal (CN) subjects from the Alzheimer’s Disease Neuroimaging Initiative (ADNI). We use a 5-layer feed-forward deep neural network (DNN) to derive relevance scores of input features and show that an empirically selected subset of features improves accuracy scores for patient classification. The common distinctive salient brain regions were in the uncus and medial temporal lobe which closely correspond with previous studies. The proposed methods have cross-modal applications with several neuropsychiatric disorders.

## 1 Introduction

Neural networks have been successfully used to classify functional neuroimaging scans of patients suffering from different stages of cognitive impairment. However, the black box nature of neural networks is a hindrance to the task of identifying salient brain regions and connections that encode differences across brain states. Such an interpretation can help characterize anomalies in the brain or differences across pathological brain states. Furthermore, knowledge of salient brain features can aid in improving the accuracy of prediction of brain states and reveal neural underpinnings of brain disease.

One can derive salient features using statistical analysis by finding differences between class-samples prior to classification. However, the presence of widespread inter-subject variability even between samples belonging to the same class, especially in the association cortices related to cognition [12], raises concerns about the precision of such methods. Therefore, recent attempts have been made to “decode” ^1^ the relevance given to input features by the classifier. Jang et al. [8] showed how trained weights in different layers of a feed-forward neural network can accurately map to four sensorimotor task states. Consequently, several works that use a gradient-based decoder with different neural network architectures have came up. Li et al. [9] used gradient-based decoding and found its principal components using temporal fMRI data to identify functional signatures in resting and task states. Similarly, Floren et al. [6] used gradient-based decoding on a feed-forward neural network to identify brain regions involved in different visual processing tasks. However, these studies do not validate their derived feature set and the existence of issues like gradient saturation and discontinuities due to bias terms need to be taken into account before using gradient-based decoders on neural networks [14].

There have been several attempts to circumvent the issues caused from gradient-based approaches, such as Integrated Gradients [15], DeepLIFT [14] and SHAP [10]. The idea is to find the contribution of each layer at the output layer and backpropagate the contributions of all the layers at the input layer. One such method, DeepLIFT, gives consideration to negative and positive contributions and computes the relevance scores very efficiently in a single pass, thus addressing the issues with gradient based approaches. In this paper, we used DeepLIFT to identify brain connections that are associated with AD and MCI. We used the ADNI dataset, the largest publicly available dataset for AD and MCI together with fMRI scans for all subjects from the same scanner.

We compared the performance of our classifier with several studies performed for classification of AD and its prodromal stage, MCI, using functional connectivity features [1,17,2,7,11,16]. By evaluating and selecting features leading to classification, we made the following novel contributions:

– Uncovered salient brain connections and regions using a reference-based decoder and show that retraining the model on an empirically relevant subset of features results in higher accuracy.
– Reported accuracies of classification for CN/MCI, MCI/AD and CN/AD, whereas previous studies performed classification only on a subset of these classes.
– Achieving state-of-the-art accuracies with the DNN in comparison to previous studies using functional connectivity features on the ADNI dataset.

## 2 Methods

### 2.1 Dataset and preprocessing

Functional and structural MRI data used in the preparation of this article were obtained from the ADNI database (`adni.loni.usc.edu`). The ADNI was launched in 2003 as a public-private partnership, led by Principal Investigator Michael W. Weiner, MD. The primary goal of ADNI has been to test whether serial magnetic resonance imaging (MRI), positron emission tomography (PET), other biological markers, and clinical and neuropsychological assessment can be combined to measure the progression of mild cognitive impairment (MCI) and early Alzheimers disease (AD). The rs-fMRI data for each subject consisted of 140 or 200 functional volumes, acquired with the following parameters: repetition time (TR) = 3000 ms; echo time (TE) = 30 ms; flip angle = 80^°^; slice thickness = 3.313 mm; and 48 slices. Results included in this manuscript come from pre-processing performed using fMRIPprep [5]. The subject details are given in table 1. As there were too few scans for MCI, we combined early MCI, MCI and late MCI into one category. Briefly, subjects classified as AD fulfilled the criteria for AD laid down by National Institute of Neurological and Communicative Disorders and Stroke and the Alzheimers Disease and Related Disorders Association. Subjects classified as MCI had a memory complaint, objective memory loss measured by education adjusted scores on Wechsler Memory Scale Logical Memory II, absence of dementia and significant levels of impairment in other cognitive domains. The CN subjects did not suffer from depression, cognitive impairment, or dementia.

**Table 1:** ADNI subject details and Mini-Mental State Examination (MMSE) and Clinical Dementia Rating (CDR) scores

|  | Cognitively Normal<br>(CN) | Mild Cognitive<br>Impairment (MCI) | Alzheimer’s Disease<br>(AD) |
| --- | --- | --- | --- |
| No. of subjects/scans | 49/196 | 90/356 | 29/103 |
| Male/Female | 22/27 | 46/44 | 15/14 |
| Age | $76.6 \pm 5.5$ | $74.4 \pm 6.7$ | $75.4 \pm 8.2$ |
| MMSE score | $28.9 \pm 1.1$ | $27.3 \pm 2.4$ | $21.2 \pm 2.6$ |
| CDR score | $0.13 \pm 0.3$ | $0.68 \pm 0.4$ | $1.13 \pm 0.6$ |

We selected 264 anatomically and functionally diverse ROIs covering the entire cerebral cortex from the Power atlas[13], and computed the mean time series of all voxels within a sphere of radius 2.5 mm around each ROI. We derive functional connectivity between the ROIs by using Pearson correlation and use them as features for the models (without thresholding).

### 2.2 Feedforward neural network

For each scan, we obtained pair (***x***, *d*) where ***x*** = (*x*_*i*_) is the input feature vector and *d* ∈ *D* is the sample label. We considered a DNN of *L* layers with *L* − 1 layers having rectified linear unit activation and a final softmax layer. Let the weights and biases of the layer *l* be given by ***W***_*l*_ and ***b***_*l*_, respectively. The output of layer *l* ≠ *L* is given by

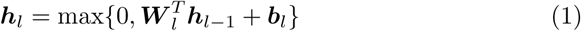

For the input layer, ***h***_0_ = ***x***. For the output softmax layer, the output is given by the probability of a sample ***x*** belonging to class *k*

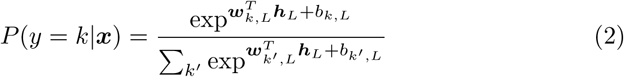

where *k ∈ {*1, …, *K}* represents class labels and the output layer weight ***W*** _*L*_ = [***w***_*k,L*_] and bias ***b***_*L*_ = (*b*_*k,L*_).

We used the cross-entropy cost to learn the parameters of the network. We performed 5-fold cross validation to select the best parameters for the DNN. We added dropouts to the hidden layers and imposed early stopping to prevent overfitting.

### 2.3 Decoding the relevance of input features

Let *f* be the neural network mapping input ***x*** to output *y*. We find a simpler *explanation model g* that is interpretable and an approximation to the model *f*. Let the number of neurons in layer *l* be *n*_*l*_. Based on an appropriate reference, let us assign to each neuron at layer *l* its contribution 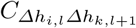 to the change in the output:

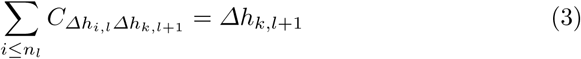

when 0 *< l < L* − 1 and

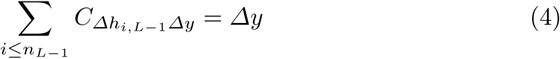

when *l* = *L* − 1, where *Δh*_*i,l*_ is the change of the neurons of layer *l* due to the input relative to the reference. It is also given that [14]:

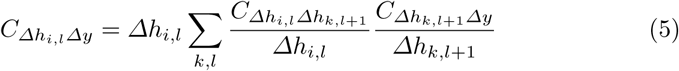

Using (3), (4) and (5), we compute the contribution of neurons in all layers 0 *≤ l < L* to the output *y* given by 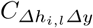 using backpropagation of contributions from output layer to the input. To compute 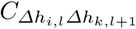, we use the Linear, Rescale and RevealCancel rule from [14]. Assuming that the reference input is given by ***x***_*r*_ and the original input by ***x***, we can substitute *Δy* = *f* (***x***) − *f* (***x***_*r*_) and *g*(***x***) = *f* (***x***), which gives us an equation for the model *g*:

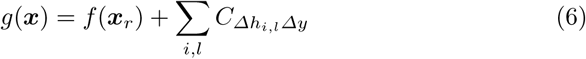

The deepLIFT method [14] computes feature relevance scores based on change in the output from a reference input. This allows information to propagate across the layers of the network even when the gradient is zero. For our binary classification problem, we computed 3 types of references, viz. sample mean, sample mode and taking a sample which has the least L2 norm from the rest of the samples. Out of these references, we empirically found the mean reference to be the most representative of the rest of the samples.

### 2.4 Recursive elimination of irrelevant features

A valid decoder can aid in selecting a subset of input features, which give an enhanced classification performance. To understand the performances of the DNN with different subset of features, we eliminated irrelevant features. We removed a percentage of features recursively and retrained the classifier on the reduced feature set. Since the number of features are different at each iteration and the DNN parameters are also updated accordingly. The threshold at which the performance of the classifier is significantly worse (*p <* 0.05) than the peak classifier performance is reported.

## 3 Results

### 3.1 Classifier performance

We performed hyperparameter tuning for the feedforward neural network and the support vector machine (SVM) model to obtain the highest classification accuracy. We varied the number of hidden layers, number of neurons in each layer, the optimizer, batch size, learning rate and learning rate decay for the feedforward network. For both linear SVM and Radial Basis Function SVM, we tried a range of parameters (*C* ∈ {0.001, 0.01, 0.1, 1, 10}, *γ* ∈ {0.001, 0.01, 0.1, 1}). We selected the best hyperparameters by performing a grid search on the average accuracy of different folds obtained from 5-fold CV.

The performance of a 5-layer DNN along with other baseline methods like SVM are shown in Table 3. All the samples were included in the test set at least once by dividing the data into 5 folds and using 1 fold for testing and the rest for training. Relevance scores for each feature were computed using the trained DNN and 10% of the features were recursively removed based on the relevance scores. We found that using a subset of salient features gave a higher accuracy, and we obtained peak accuracies for 28.3% features for CN/MCI, for 59% features for MCI/AD and for 43% features for CN/AD for the 5-layer feedforward network shown in Table 2. For all cases, the classifier gave a significantly worse accuracy than the peak accuracy when less than 25% of the features were used.

**Table 2:** Parameters of the 5 layer feedforward network giving the highest accuracy

| Parameter | CN/MCI | MCI/AD | CN/AD |
| --- | --- | --- | --- |
| Input Nodes | 9825 | 20482 | 14927 |
| Hidden Nodes (each layer) | 1415-283-6 | 2950-590-295 | 2150-86-9 |
| Learning Rate | 0.002 | 0.0012 | 0.0005 |
| Batch size | 4 | 2 | 4 |
| Optimizer | SGD | SGD | SGD |
| Decay | 0.001 | 0.001 | 0.0001 |

**Table 3:** Comparison of functional connectivity based classifiers

| Model | CN/MCI | MCI/AD | CN/AD |
| --- | --- | --- | --- |
| 3 layer DNN (all features) | 85.1% ± 3.0% | 91.9% ± 2.9% | 91.0% ± 4.9% |
| 3 layer DNN (feature subset) | **86.4%** ± **3.4%** | **92.4%** ± **3.2%** | **92.0%** ± **4.3%** |
| Cutoff ($p < 0.05$) | 20.6% | 22.8% | 15.0% |
| Linear SVM ($C = 0.01$) | 83.2% ± 2.9% | 85.9% ± 3.2% | 89.7% ± 4.1% |
| RBF SVM ($C = 10, \gamma = 0.001$) | 81.6% ± 2.0% | 88.6% ± 2.4% | 87.7% ± 3.6% |
| Challis et al. (In-house) [1] | 75% | **90%** | - |
| Zhang X. et al. (ADNI) [17] | **87.5%** | - | - |
| Chen X. et al. (ADNI) [2] | 78.7% | - | - |
| Guo H. et al. (In-house) [7] | - | - | **91.6%** |
| Meszlenyi et al. (CoRR) <sup>2</sup>[11] | 72.9% | - | - |
| Yu. et al (ADNI) [16] | 84.8% | - | - |

We achieved state-of-the-art accuracies for MCI/AD and CN/AD classification in comparison with studies that used functional connectivity as features. Zhang X. et al. [17] had a marginally better accuracy for CN/MCI, which could be explained by their usage of only MCI subjects vs CN, while we combined early MCI with MCI which resulted in a more difficult classification task. The minor difference in CN/AD results can be due to the different dataset used in Guo H. et al. [7].

### 3.2 Salient features found by the decoder

Using the relevance scores of functional connections, we compute the importance of brain ROI by summing up functional connections incident on them. We plot the ROIs with top 10% relevance score using Nilearn (Fig. 1).

**Fig. 1:**
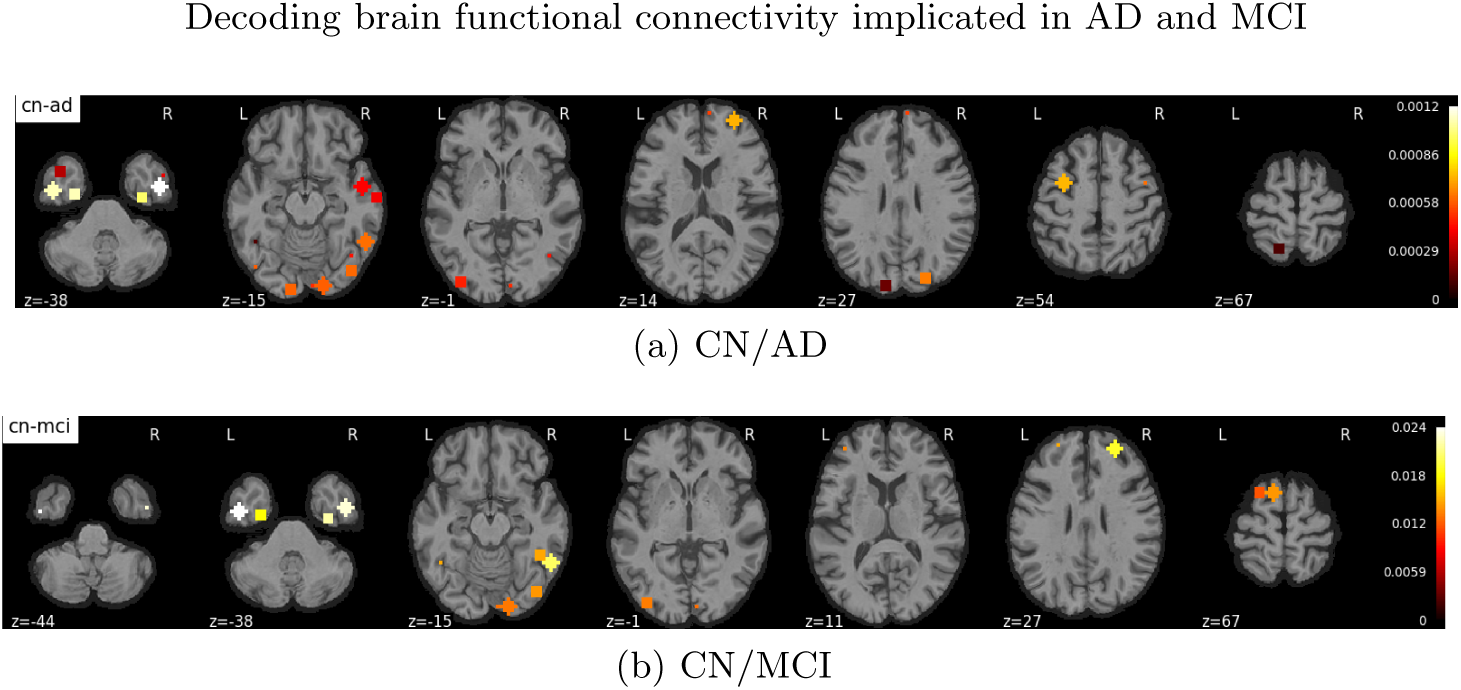
The axial top view of ROI with top 10% relevance scores discriminating AD and MCI patients from CN subjects.

Besides observing the position of important ROIs in the brain, it is also of interest to understand the anatomical locations of functional connections between brain ROIs that are important for classification (Fig. 2). However, since the anatomical labels for the ROIs from the Power Atlas are absent, we use the anatomical labels of ROIs from the Crossley atlas [3] and map Power ROI to Crossley ROI by computing the Euclidean distance between the ROIs from the two atlases. Across MCI/AD, CN/AD and CN/MCI, we found two regions in the uncus, an anterior extremity of the parahippocampal gyrus, to be highly relevant (Fig. 1 and 2), which is consistent with early atrophy observed in the area. As reported before [4], we found several medial temporal lobe regions in the parahippocampus, fusiform, lingual and angular gyrus to be highly discriminative for both AD and MCI. We found regions in the superior frontal gyrus (containing both medial and lateral regions) and temporal poles to be highly relevant specifically for MCI patients, which has been reported before [11,7].

**Fig. 2:**
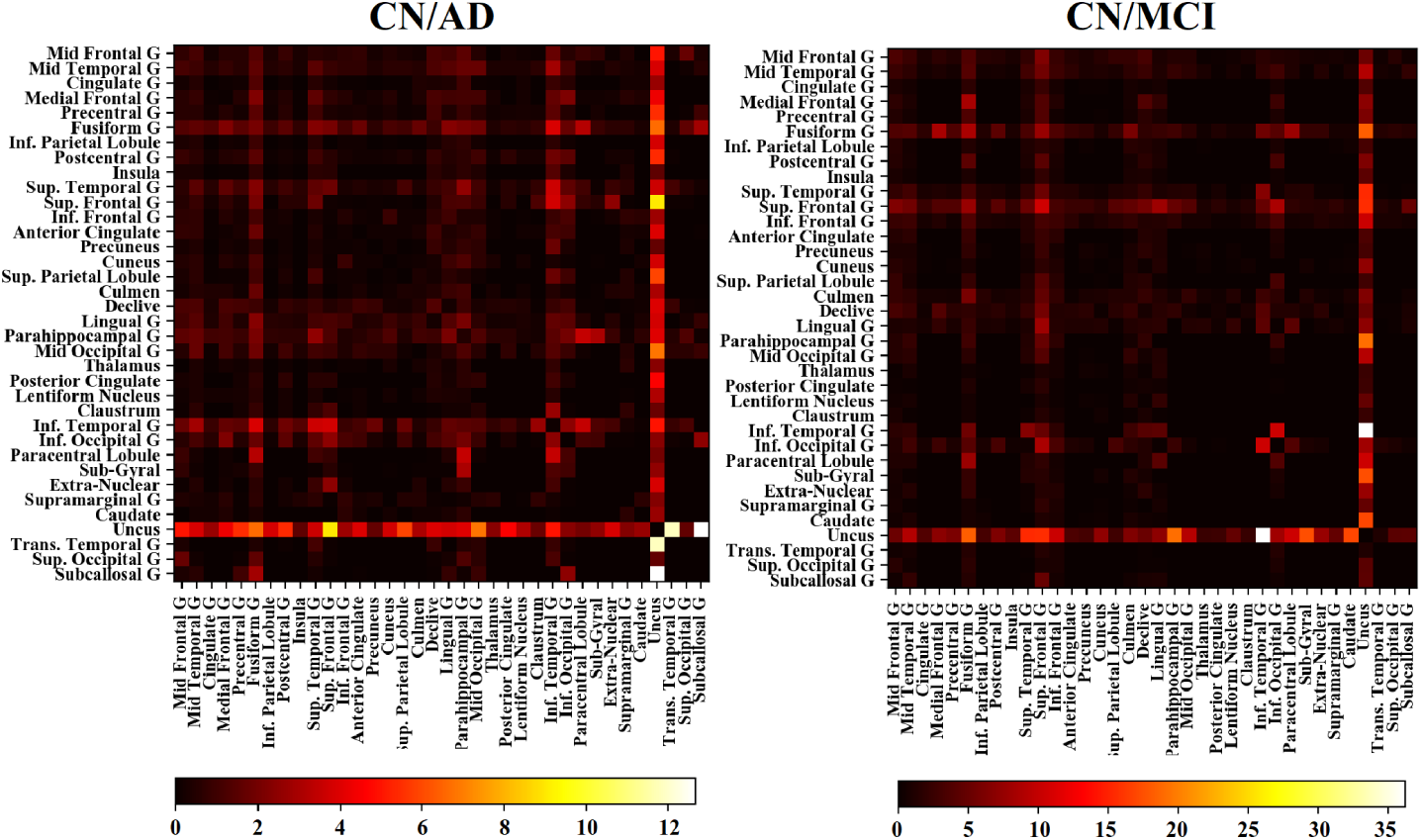
The figure shows the relevance scores of functional connections between brain regions derived from the Power atlas while differentiating MCI and AD patients from CN subjects. Mid: Middle, Inf: Inferior, Sup: Superior, G: Gyrus

## 4 Conclusion

We proposed an approach of recursively eliminating features with low relevance scores so as to obtain a leaner DNN. This is a critical issue since fMRI data often has many more features than data instances to train the DNN. We used a 5-layer feedforward DNN to classify fMRI scans of CN subjects and patients suffering from MCI and AD from the ADNI dataset. Using feature relevance scores from a reference based decoder, we showed that using a subset of important input features indeed improves the classification of fMRI brain scans. We achieved state-of-the-art classification accuracy for classification of MCI/AD and CN/AD classification and comparable accuracies for CN/MCI classification. We further studied the brain regions identified as important by the decoder and found these to be supported by previous studies. The proposed methods can be used for biomarker detection for several neurological ailments and subsequently aid in better computer-aided-detection.

## 5 Acknowledgement

This work was partially supported by AcRF Tier 1 grant RG 19/15 of Ministry of Education, Singapore. Data collection and sharing for this project was funded by the Alzheimer’s Disease Neuroimaging Initiative (ADNI) (National Institutes of Health Grant U01 AG024904) and DOD ADNI (Department of Defense award number W81XWH-12-2-0012). ADNI is funded by the National Institute on Aging, the National Institute of Biomedical Imaging and Bioengineering, and through generous contributions from the following: AbbVie, Alzheimers Association; Alzheimers Drug Discovery Foundation; Araclon Biotech; BioClinica, Inc.; Biogen; Bristol-Myers Squibb Company; CereSpir, Inc.; Cogstate; Eisai Inc.; Elan Pharmaceuticals, Inc.; Eli Lilly and Company; EuroImmun; F. Hoffmann-La Roche Ltd and its affiliated company Genentech, Inc.; Fujirebio; GE Healthcare; IXICO Ltd.; Janssen Alzheimer Immunotherapy Research & Development, LLC.; Johnson & Johnson Pharmaceutical Research & Development LLC.; Lumosity; Lundbeck; Merck & Co., Inc.; Meso Scale Diagnostics, LLC.; NeuroRx Research; Neurotrack Technologies; Novartis Pharmaceuticals Corporation; Pfizer Inc.; Piramal Imaging; Servier; Takeda Pharmaceutical Company; and Transition Therapeutics. The Canadian Institutes of Health Research is providing funds to support ADNI clinical sites in Canada. Private sector contributions are facilitated by the Foundation for the National Institutes of Health (www.fnih.org). The grantee organization is the Northern California Institute for Research and Education, and the study is coordinated by the Alzheimers Therapeutic Research Institute at the University of Southern California. ADNI data are disseminated by the Laboratory for Neuro Imaging at the University of Southern California.

## Footnotes

1 We refer to deriving the relevance scores of features as decoding, which is different from the decoding done in autoencoders.

2 Consortium for Reliability and Reproducibility

## Notes

http://adni.loni.usc.edu/

