## Appendix for "Decoding brain functional connectivity implicated in AD and MCI"

School of Computer Science and Engineering,
Nanyang Technological University, Singapore.
{sukrit001, yihao001, asjagath}@ntu.edu.sg

* Data used in preparation of this article were obtained from the ADNI database (adni.loni.usc.edu).

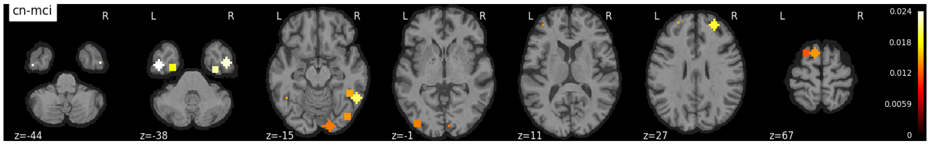


(a)

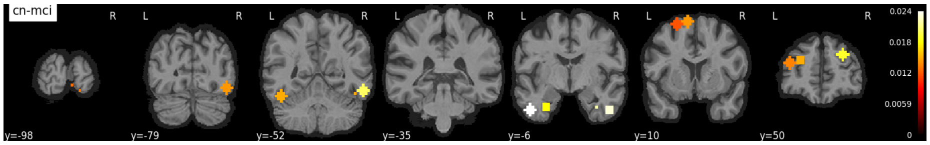


(b)

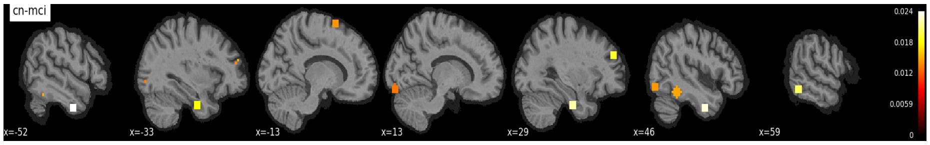


(c)

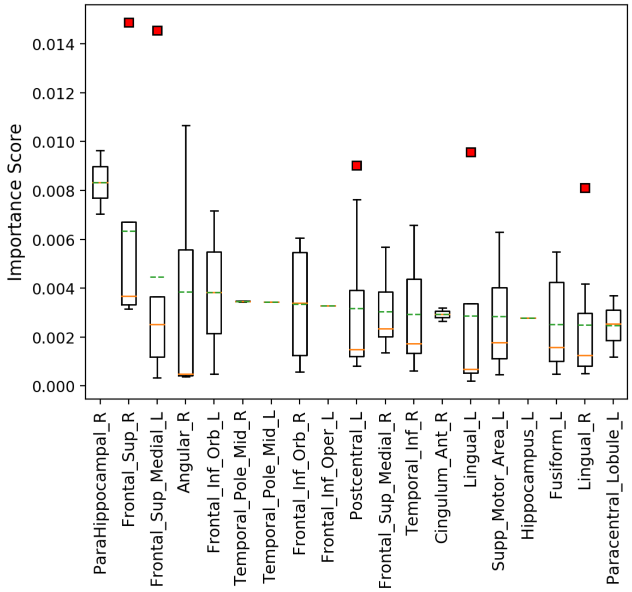


(d)

Fig. 1: Anatomical locations and labels of salient ROI differentiating MCI patients from CN subjects. (a), (b) and (c) are the axial, coronal and sagittal views of top 10% ROI derived from the remaining features after recursive elimination; and (d) are the anatomical location labels (in decreasing order of salience) for brain regions which differentiate MCI patients from CN.

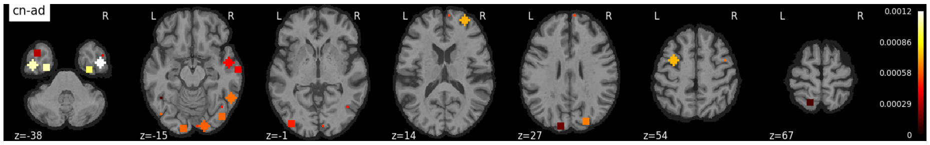


(a)

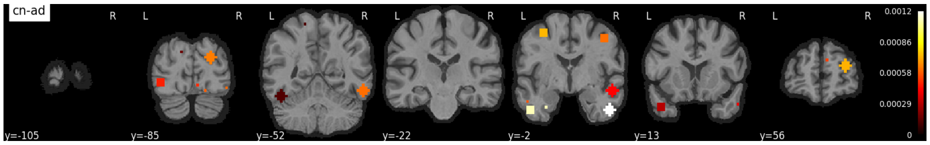


(b)

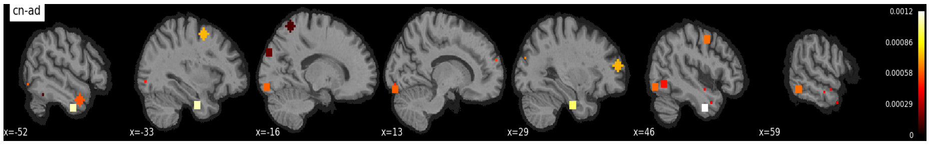


(c)

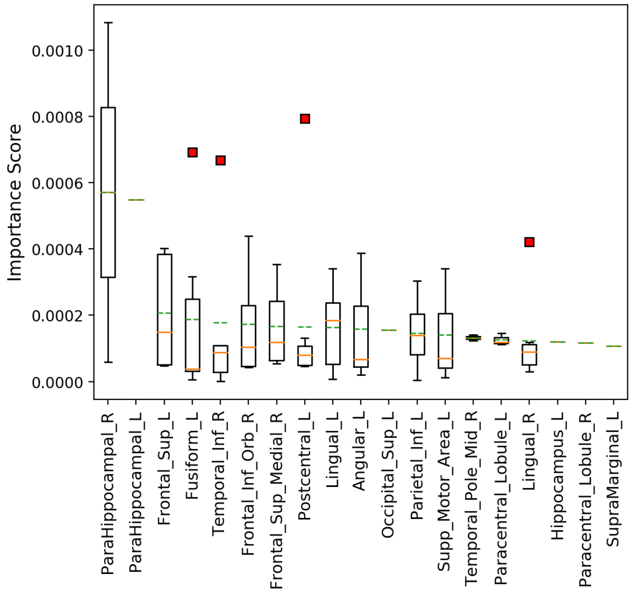


(d)

Fig. 2: Anatomical locations and labels of salient ROI differentiating AD patients from CN subjects. (a), (b) and (c) are the axial, coronal and sagittal views of top 10% ROI derived from the remaining features after recursive elimination; and (d) are the anatomical location labels (in decreasing order of salience) for brain regions which differentiate AD patients from CN.

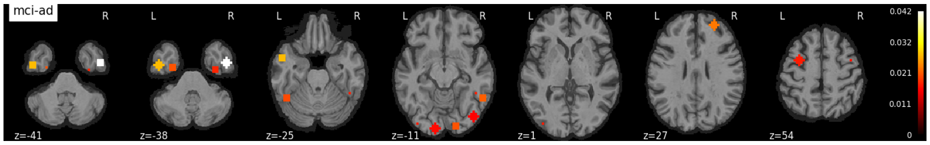


(a)

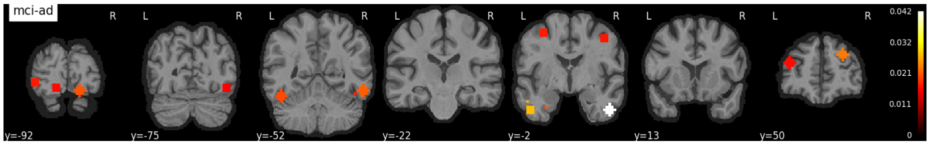


(b)

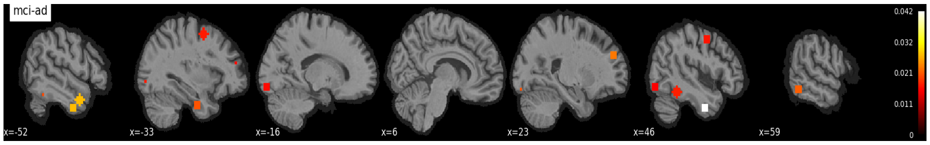


(c)

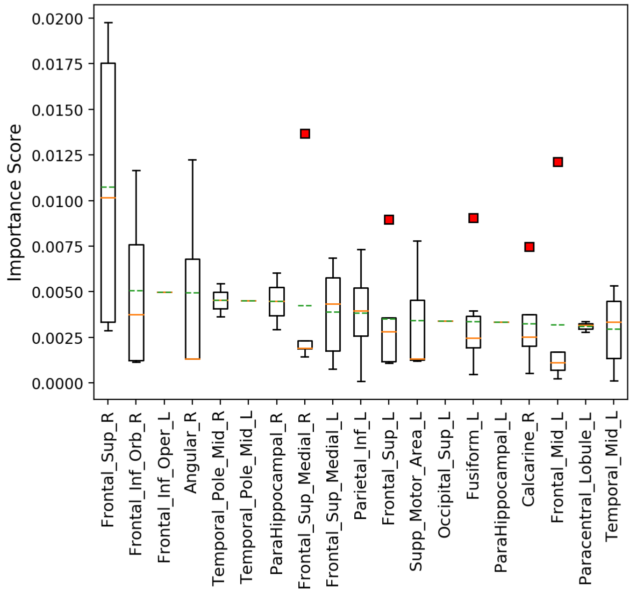


(d)

Fig. 3: Anatomical locations and labels of salient ROI differentiating MCI patients from AD patients. (a), (b) and (c) are the axial, coronal and sagittal views of top 10% ROI derived from the remaining features after recursive elimination; and (d) are the anatomical location labels (in decreasing order of salience) for brain regions which differentiate MCI patients from AD patients.

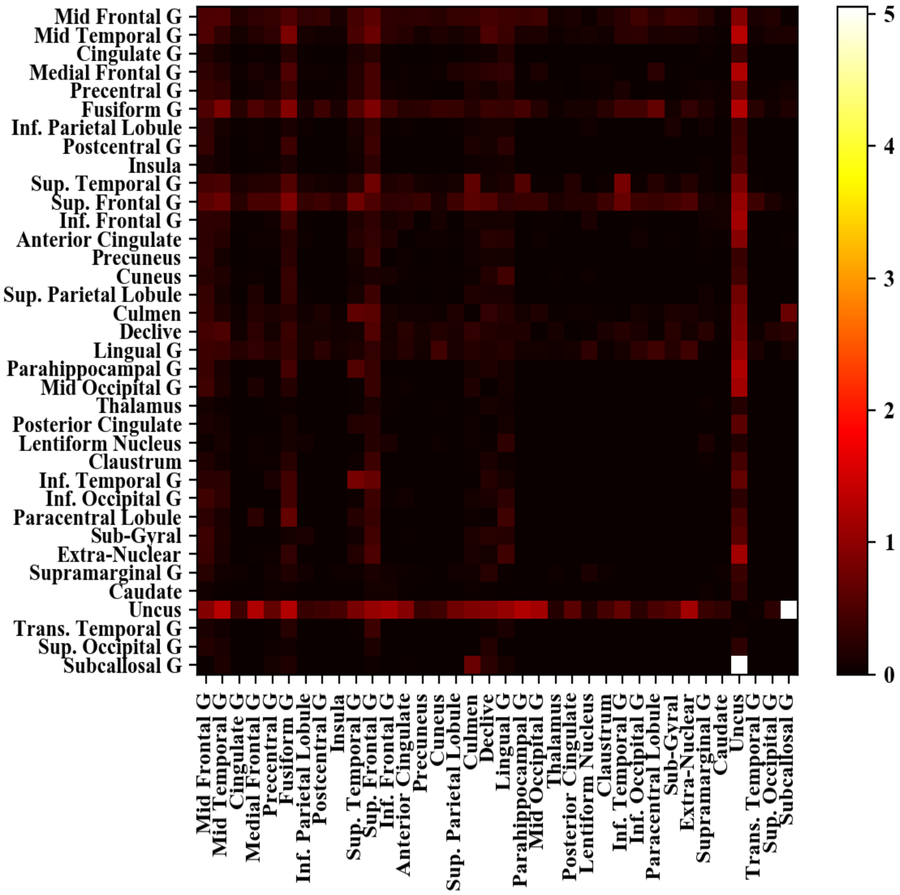


Fig. 4: The salient functional connections between different brain anatomical locations which differentiate MCI and AD patients.
